# Scalable Design of paired CRISPR Guide RNAs for Genomic Deletion

**DOI:** 10.1101/052795

**Authors:** Carlos Pulido-Quetglas, Estel Aparicio-Prat, Carme Arnan, Taisia Polidori, Toni Hermoso, Emilio Palumbo, Julia Ponomarenko, Roderic Guigo, Rory Johnson

## Abstract

Using CRISPR/Cas9, diverse genomic elements may be studied in their endogenous context. Pairs of single guide RNAs (sgRNAs) are used to delete regulatory elements and small RNA genes, while longer RNAs can be silenced through promoter deletion. We here present CRISPETa, a bioinformatic pipeline for flexible and scalable paired sgRNA design based on an empirical scoring model. Multiple sgRNA pairs are returned for each target. Any number of targets can be analyzed in parallel, making CRISPETa equally appropriate for studies of individual elements, or complex library screens. Fast run-times are achieved using a precomputed off-target database. sgRNA pair designs are output in a convenient format for visualisation and oligonucleotide ordering. We present a series of pre-designed, high-coverage library designs for entire classes of non-coding elements in human, mouse, zebrafish, *Drosophila* and *C. elegans*. Using an improved version of the DECKO deletion vector, together with a quantitative deletion assay, we test CRISPETa designs by deleting an enhancer and exonic fragment of the *MALAT1* oncogene. These achieve efficiencies of ≥50%, resulting in production of mutant RNA. CRISPETa will be useful for researchers seeking to harness CRISPR for targeted genomic deletion, in a variety of model organisms, from single-target to high-throughput scales.

## Introduction

CRISPR/Cas9 is a simple and versatile method for genome editing that can be applied to deleting virtually any genomic region for loss-of-function studies. Recent vector tools have been developed for complex library cloning that are compatible with pooled screening (1,2). Whether performing pooled screens on hundreds of targets, or deletion of a single target, researchers need to design efficacious pairs of sgRNAs. We present here a flexible and scalable software pipeline to address the needs of both types of project.

CRISPR/Cas9 makes it possible to investigate the function of genomic elements in their endogenous genetic context. The Cas9 nuclease is recruited to desired genomic sites through its binding to an engineered, single guide RNA (sgRNA) (3). Early studies focussed on protein coding genes, utilizing individual sgRNAs to induce small indel mutations in genomic regions encoding target proteins’; open reading frame (ORFs). Such mutations frequently give rise to inactivating frameshift mutations, resulting in complete loss of function (4,5). The delivery of a single sgRNA in such experiments is technically straightforward, and can be scaled to genome-wide, virally-delivered screens.

CRISPR has also been brought to bear on non-coding genomic elements, including regulatory regions and non-coding RNAs, which have traditionally resisted standard RNA interference (RNAi) (6,7). Unlike coding genes, functional knockout of non-coding elements with a single sgRNA is probably not practical, because small indel mutations caused by single sgRNAs are less likely to ablate function. Instead, a deletion strategy has been pursued: a pair of sgRNAs are used to recruit Cas9 to sites flanking the target region (1,7). Simultaneous strand breaks are induced, and non-homologous end joining (NHEJ) activity repairs the lesion. In a certain fraction of cases, this results in a genomic deletion with a well-defined junction.

Cas9 targeting is achieved by engineering the 5’ region of the sgRNA. This hybridises to a complementary “protospacer” region in DNA, immediately upstream of the “protospacer adjacent motif” (PAM) (8). For the most commonly used *S. pyogenes* Cas9 variant, the PAM sequence consists of “NGG”. A growing number of software tools are available for the selection of optimal protospacer targeting sequences (9-15). The key selection criteria are (1) the efficiency of a given sequence in terms of generating mutations, and (2) “off-targeting”, or the propensity for recognising similar, yet undesired, sites in the genome. Based on experimental data, scoring models for on-target efficiency have been developed, for example that presented by Doench et al (13). At the same time, tools have become available for identifying unique sgRNA sites genome-wide, mitigating to some extent the problem of off-targeting (16). However, few tools presented so far are designed for large-scale designs, and to the best of our knowledge, none was created to identify optimal sgRNA *pairs* required for deletion studies.

To address this need, we here present a new software pipeline called CRISPETa (CRISPR Paired Excision Tool) that selects optimal sgRNAs for deletion of user-defined target sites. The pipeline has two useful features: first, it can be used for any number of targets in a single, rapid analysis; second, it returns multiple, optimal pairs of sgRNAs, with maximal predicted efficiency and minimal off-target activity. The pipeline is available as both standalone software and as a user-friendly webserver. In addition, we make available a number of pre-designed deletion libraries for various classes of non-coding genomic elements in a variety of species. Finally, we validate CRISPETa predictions experimentally by means of an improved version of the published DECKO deletion technique (1). Using a quantitative deletion assay, we find that CRISPETa predictions are highly efficient in deleting fragments of a human gene locus’; resulting in detectable changes to the cellular transcriptome. CRISPETa is available at www.crispeta.crg.eu.

## Materials and Methods

### Details of CRISPETa Code

The pipeline is outlined in Figure 1A. As input, CRISPETa requires a standard BED6-format file describing all target regions. This file must contain coordinates of one or more targets. Unstranded entries are assigned to the + strand, while those without identifiers are assigned a random ID. CRISPETa first defines design regions based on parameters *g/du/dd/eu/ed* (see Table 1 for full list of parameters) (Figure 1A,B), and extracts their sequences using the BEDtools *getfasta* function. Design regions are searched for canonical PAM elements (NGG) using a regular expression. For every such PAM, a total of 30 nucleotides (NNNN[20nt]NGGNNN) are stored. Protospacers containing the RNA Pol III stop sequence (TTTT) are removed.

**Figure 1:**
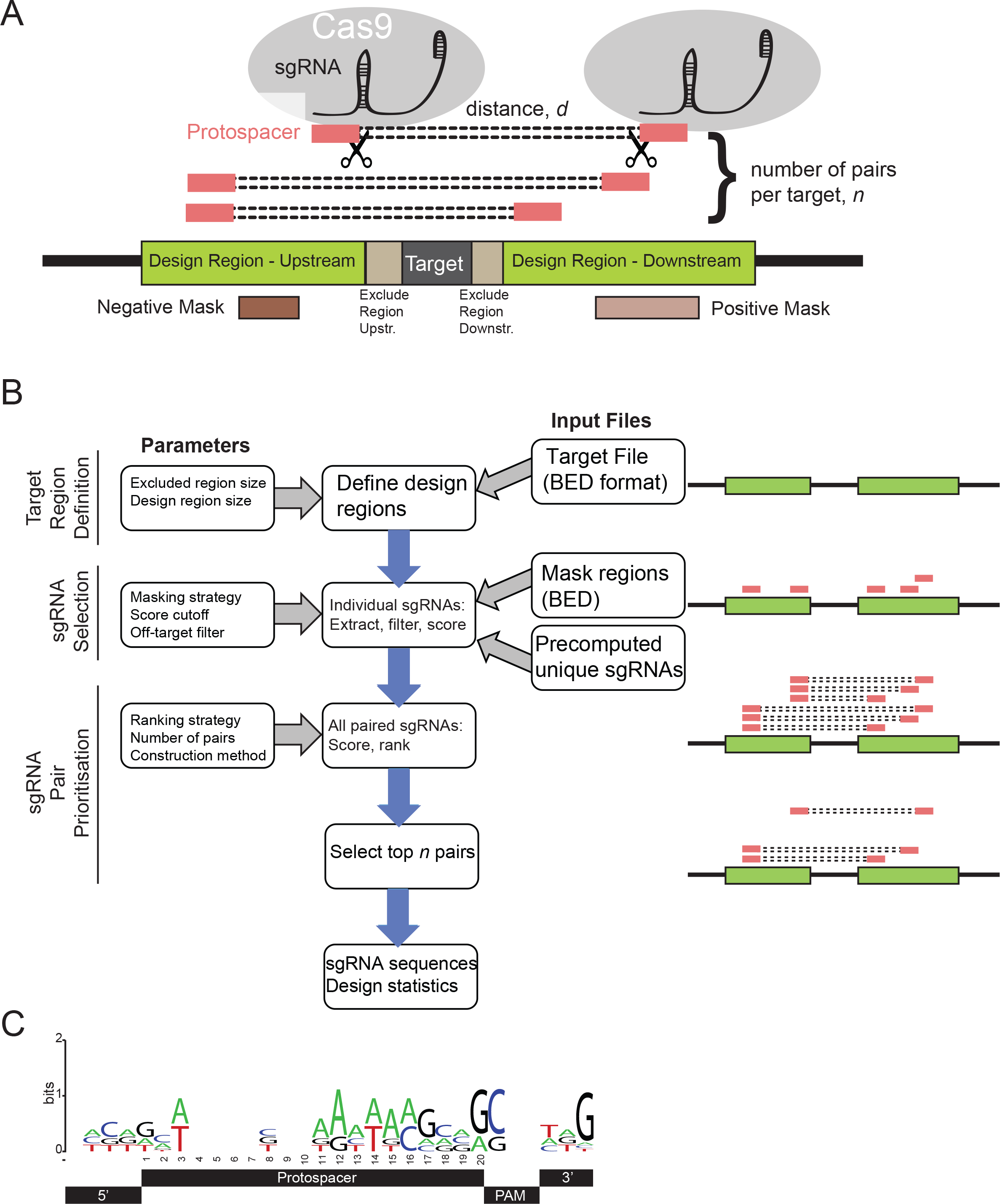
Overview of CRISPETa pipeline. (A) Schematic of CRISPR-mediated genomic deletion. The aim is elimination of the Target region through recruitment of a pair of Cas9 proteins. Red boxes represent protospacers, the 20 bp upstream of a PAM and recognised by the sgRNA. (B) The CRISPETa workflow. (C) Sequence preferences that drive the efficiency prediction score. Note that positions outside the protospacer are also predictive of efficiency.

**Table 1.**
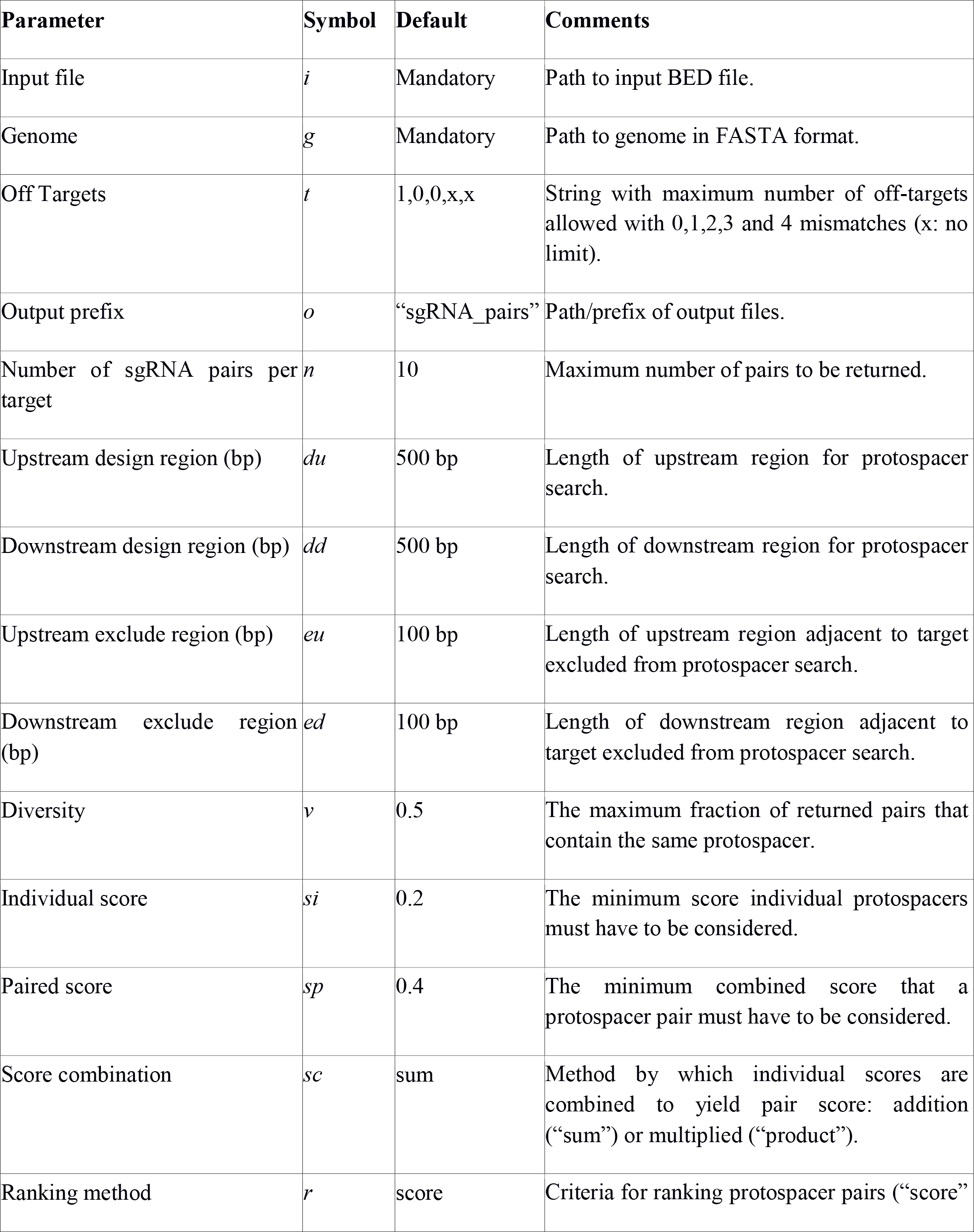
User-defined parameters.

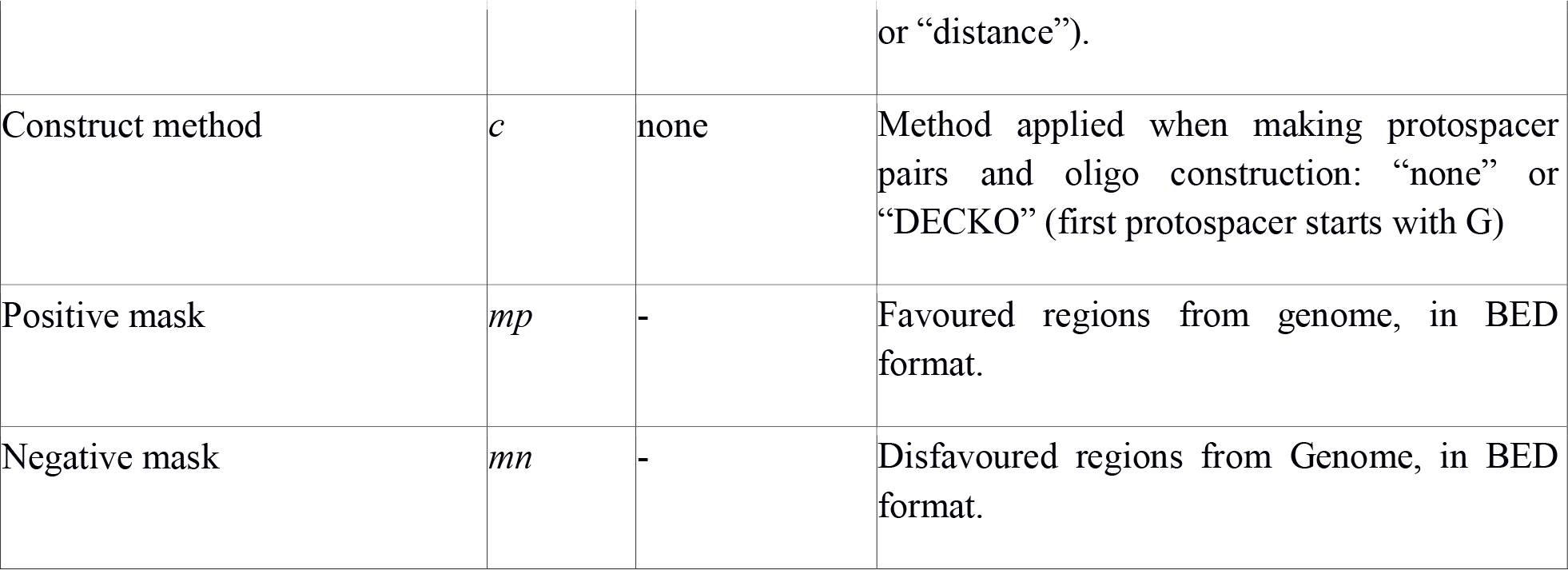

Next, candidate protospacers are searched against a precomputed, database-stored list of potential protospacers and their number of similar sequences with up to 4 mismatches, genome-wide (see “Off-target analysis” section, below). By default, protospacers with one or more off-targets with ≤2 mismatch are discarded (this cutoff can be modified by the user through parameter *t*). Remaining protospacers are then compared with the positive and negative mask BED files using BEDtools *intersectBed.* Candidate sequences not fully overlapping the positive mask file, or overlapping the negative mask by one basepair, are tagged as “disfavoured”. Next, 30mer regions encompassing remaining protospacers, including disfavoured ones, are assigned an efficiency score (see below) between 0 and 1, and those above the score threshold (controlled by parameter *si*) are carried forward.

Next, candidate sequences are assembled into pairs and filtered. For each target region, all possible pairs of upstream and downstream candidates are generated. If pairs are designed for DECKO cloning (which utilizes the U6 promoter for the 5’ sgRNA gene, controlled by c), an additional filter is applied: sgRNA pairs are rearranged as necessary, such that all output pairs have the first sgRNA starting with G, and any pairs where neither commences with G are removed. A combined score for the resulting pairs is computed. By default, this is the sum of the two individual sgRNA scores, but users may choose to define the pair score as the product of individual scores (parameter *sc*). Pairs are now filtered with a pair score threshold, and ranked by score (or, optionally, reversed rank by distance, parameter *r*). An optional “diversity” cutoff can be used to remove pairs such that no individual candidate sequence appears in more than a given fraction of returned pairs (parameter *ν*). Finally the program returns the top ranked pairs up to the maximum number specified by the user, *n*.

CRISPETa is implemented in Python and available for download from git-hub and the CRISPETa web-server (see availability below).

### Target features and mask files

All target sets and mask files were prepared in BED format, and obtained in April 2016. Coding genes were obtained from the Gencode v19 annotation, filtered for the “protein_coding” biotype (17). CTCF binding sites for GM12878 cells were downloaded from ENCODE data hosted in the UCSC Browser (18). Enhancers were obtained from Vista (19). Pre-miRNAs were obtained from miRBASE (20). Disease-associated SNPs were obtained from the GWAS database (http://www.ebi.ac.uk/gwas/api/search/downloads/full). Ultraconserved regions were obtained from UCNEbase (21). For human positive and negative masks we used DNaseI hypersensitive sites identified through genome-wide profiling in 125 diverse cell and tissue types by the ENCODE consortium (22) and RepeatMasker repetitive regions (23)’ respectively. To generate random intergenic locations’; the entire span of all Gencode v19 genes (both coding and noncoding’ introns and exons)’ in addition to 100 kb up-and downstream’ were subtracted. Random locations were selected within the remaining regions.

### Off-target analysis

Off-target analysis was performed using Crispr-Analyser (16). We searched for all canonical PAM regions (NGG) in the genome and stored the 20nt that precedes each. Then using “search” and “align” options we obtained the number of off-targets with 0,1,2,3 and 4 mismatches for each unique 20mer. This data was stored in a MySQL database. Precomputed files containing this information for various genomes can be directly downloaded (see **CRISPETa availability** section). Downloadable files contain 6 comma-separated fields in this order: sequence of the sgRNA without the PAM sequence and the number of off-targets with 0,1,2,3 and 4 mismatches for this sgRNA. These files can be used as input for CRISPETa-MySQL module to generate the MySQL database.

### CRISPETa availability and webserver

CRISPETa can be run through the web-server (http://crispeta.crg.eu) or locally. The software runs on python2.7. In order to run CRISPETa locally two additional programs are required: BEDtools and MySQL. Source code to run locally can be found on git-hub (https://github.com/guigolab/CRISPETA) and also on “Get CRISPETa” section of the web-server. Source code consist of two scripts: CRISPETA.py that execute the main pipeline described above’ and crispeta_mysql.py that helps users to create the off-target MySQL database. Two other files can be found within the source code: func.py that contains all functions necessary to execute the two main scrips’ and config.py that stores the information needed to login to MySQL.

### sgRNA scoring algorithm

CRISPETa uses the scoring method developed by Doench et al (13)’ based on an experimentally trained logistic regression model employing 72 sequence features. The code was downloaded from http://www.broadinstitute.org/rnai/public/analysis-tools/sgrna-design-v1.

### Benchmarking

A test target set contains 1000 random elements from each of the individual target annotations’; for a total of 7000. Benchmarking analyses were run on a workstation running CentOS6’ 86.6 Gb of memory and 12 CPUs (Intel(R) Xeon(R) CPU E5649 @ 2.53GHz).

### DECKO2 design and molecular cloning

A detailed protocol for DECKO2 molecular cloning is available from Supplementary File 1. Selected sgRNA pairs were converted to overlapping series of 6 oligonucleotides (Figure 4B’ Supplementary File 5) using a custom design spreadsheet (available as Supplementary File 2). Note that Oligos 3&4 do not vary between experiments. Oligos were synthesised commercially and combined at a final concentration of 0.1μm’ together with 100-200ng of BsmbI-digested backbone pDECKO_mCherry in 10 μl volume’ and 10 μl of 2x Gibson mix. The latter was prepared in house’ according to the protocol described previously (1). We incubated the mixture at 50°C for 1 hour’ and fast-transformed 2 μl of this into 50 μl of z-Stbl3 competent cells (prepared with Mix and Go *E. coli* transformation kit from Zymo Research’ Cat. T3001). Resulting “intermediate” plasmids (corresponding to Figure 4B) were amplified and purified.

The Insert-2 fragment (Figure 4C) was amplified from plasmid pDECKO _GFP (1) (Addgene ID XXX) using primers “Scaffold F” and “H1 R” (Supplementary File 4)’ and gel-purified. This was inserted’ by Gibson assembly’ into the intermediate plasmid that had previously been linearized by BsmbI-digestion and column-purified. We sequence-verified these constructs with primers “Seq 3 F/R” (Supplementary File 4).

### Genotyping by PCR

gDNA was extracted with GeneJET Genomic DNA Purification Kit (Thermo Scientific). PCR was performed with primers flanking the deleted region (primers “out F/R” in Supplementary File 4).

### QC-PCR assay

gDNA was extracted with GeneJET Genomic DNA Purification Kit (Thermo Scientific) and quantitative real time PCR (qPCR) from 1.6 ng of purified gDNA was performed on a LightCycler 480 Real-Time PCR System (Roche). Primer sequences can be found in Table S3 and S4 of Supplementary File 4. Target sequence primers (Enhancer in F / Enhancer out R for enhancer’ Exon in F / Exon out R for exon) were normalised to primers GAPDH F/R amplifying a distal’ non-targeted region. Another non-targeting primer set’ LdhA F/R were treated in the same way. Data were normalised using the ΔΔCt method (24)’ incorporating primer efficiencies. The latter were estimated using a dilution series of gDNA’ and efficiency calculated by the slope of the linear region only (Supplementary File 3). We noted a decrease in efficiency at high template densities.

### Cell culture and DECKO2 knockouts

HEK293T were grown in Dulbecco’;s modified Eagle’;s medium (DMEM’ Life Technologies) and IMR90 in Eagle’;s Minimum Essential Medium (EMEM’ ATCC). Media were supplemented with 10% fetal bovine serum (FBS’ Gibco)’ 5% Penicillin-Streptomycin Streptomycin (Life Technologies). Cells were maintained at 37°C in a humid atmosphere containing 5% CO2 and 95% air. Cells were transfected with Lipofectamine 2000 (Life Technologies) following the manufacturer’;s protocol. For lentivirus production, pDECKO_mCherry plasmids was co-transfected into HEK293T cells with the packaging plasmids pVsVg (Addgene 8484) and psPAX2 (Addgene 12260).

To create Cas9 stably-expressing cells, we transfected Cas9 plasmids and selected for more than 5 days with blasticidin at 10μg/ml.

### Genomic deletion at low multiplicity of infection

For lentivirus production, pDECKO_mCherry plasmids (pDECKO_mCherry_TFRC_B) (3 μg) were cotransfected into HEK293T cells with the packaging plasmids pVsVG (Addgene 8484) (2.25 μg) and psPAX (Addgene 12260) (750 ng) in 10 cm dishes. Viral supernatant was collected after 48 hours and filtered through 0.45 μm cellulose acetate syringe filter. We used between 0.5 and 1 ml of viral supernatant, along with polybrene at a final concentration of 10 μg/ml, to perform overnight infection of IMR90-Cas9BFP cells seeded approximately at 60% confluence in 6 well plates. Media was changed the following day, and half of the cells were analyzed by flow cytometry to ascertain infection rate. Remaining cells were double selected with puromycin and blasticidin for 14 days. Cells were lysed with 50 μl of Lysis Buffer (25 nM NaOH, 0.2 mM EDTA) and heated at 95°C for 30 min (25). The reaction was inactivated with Tris Buffer (40 mM Tris-HCl) and lysates centrifuged for 5 min at 4000 rpm. For genotyping, we performed qPCR directly from cell lysates with primers “TFRC_B out F” and “TFRC_B in R” and normalized against “GAPDH F/R” (see Supplementary File 4).

## Results

### The CRISPETa pipeline for paired sgRNA design

We and others have recently demonstrated the cloning of paired CRISPR targeting constructs for deletion of genomic regions. This creates the need for a design pipeline to select optimal pairs of sgRNAs. Our solution is the CRISPETa pipeline, whose principal steps are shown in Figure 1B. The guiding principles of CRISPETa are flexibility and scalability: the user has control over all aspects of the design process if desired (otherwise reasonable defaults are provided), and the design may be carried out on individual targets, or target libraries of essentially unlimited size. The full set of user-defined variables, and their default values, are shown in Table 1.

We use here the standard term “protospacer” to designate the 20 bp of genomic DNA sequence preceding the PAM sequence (8), as distinct from the sgRNA sequence itself, composed of the protospacer sequence and the constant, scaffold region.

The CRISPETa workflow may be divided into three main steps: target region definition, protospacer selection, and sgRNA pair prioritisation (Figure 1B). Given a genomic target region or regions in BED format, CRISPETa first establishes pairs of “design regions” of defined length in which to search. Design regions may be separated from the target itself by “exclude regions”. The user may also specify “mask regions”: sgRNAs falling within the positive mask are prioritised, whereas those within the negative mask will be de-prioritized (although not removed altogether). Positive masks might include regions of DNaseI-accessible chromatin, while negative masks may be composed of, for example, repetitive regions or compact chromatin.

Using this information, the entire set of potential protospacers is defined. First, the design region sequence is extracted and searched for all possible 20mer sites followed by canonical *S. pyogenes* “NGG” PAM sites-candidate protospacers. These are considered with respect to two core metrics: their potential for off-target binding, and their predicted efficiency. Off-targeting, or the number of identical or similar sites with a given number of mismatches, is estimated using precomputed data for each genome. This strategy increases the speed of CRISPETa dramatically. We created off-target databases for five commonly-studied species, human, mouse, zebrafish, *Drosophila melanogaster, Caenorhabditis elegans* (Table 2), varying widely in genome size (Figure 2A). The default off-targeting cutoff is set at (0:1, 1:0, 2:0, 3:x, 4:x), that is, sequences having no other genomic site with ≤2 mismatches. At this default, 77% of candidate protospacers are discarded in human, compared to just 13% in *Drosophila,* reflecting the relative uniqueness and compactness of the latter (Figure 2B).

**Figure 2:**
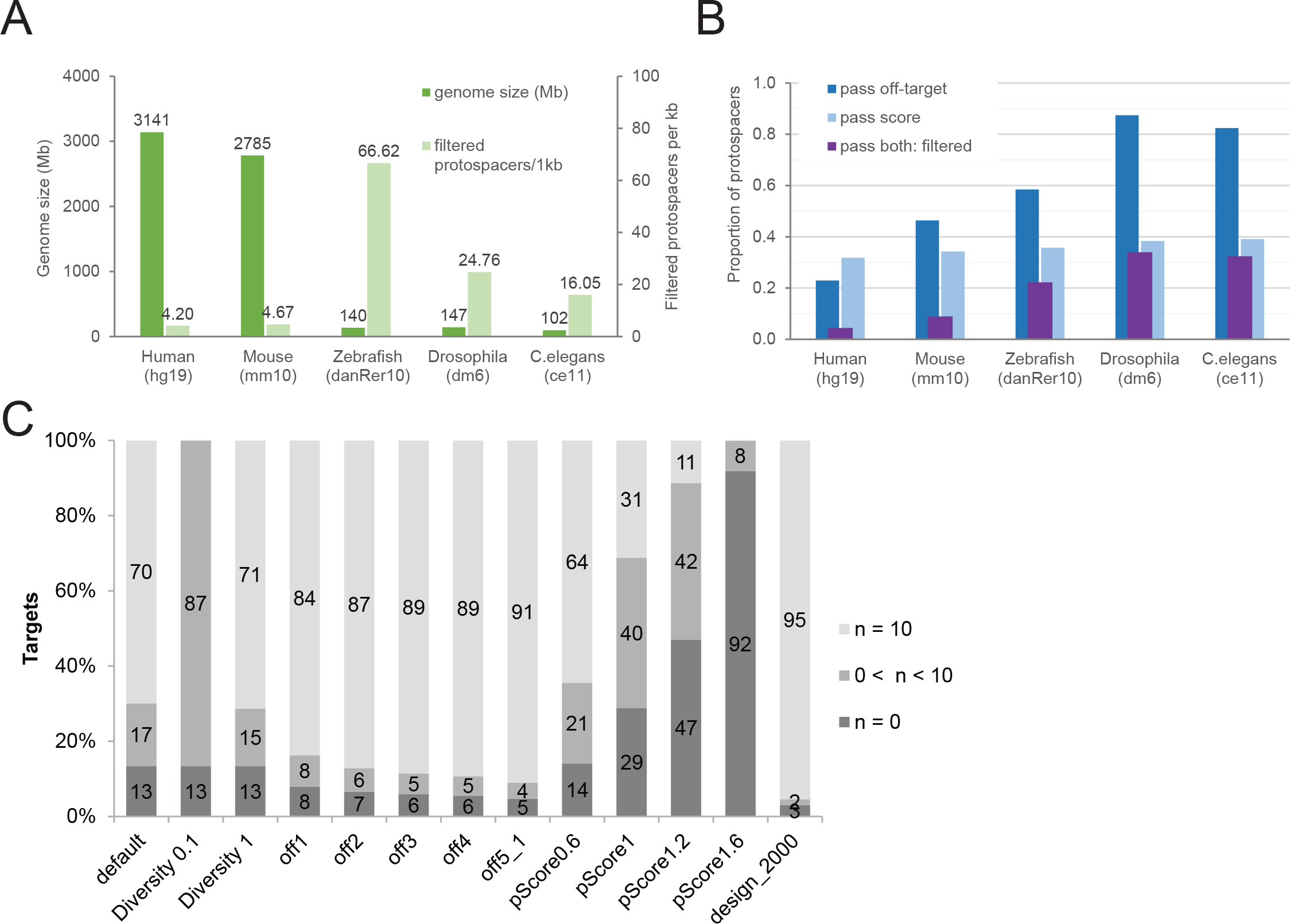
Benchmarking and performance. (A) Genome size and filtered protospacer density for the five species tested. (B) The fraction of protospacers passing filters of off-targeting, efficiency score, and both. The latter are defined as “filtered protospacers”, whose density is shown in (A). Data are displayed as a proportion of the total number of canonical PAM sequences in each genome. (C) The effect on library quality of modifying design variables. Y-axis denotes the percent of target regions, divided by: “successful”, where n=10 distinct sgRNA pair designs are returned per target; “intermediate” designs, where 0<n<10 pairs are returned; “failed” designs, where n=0 pairs are returned. CRISPETa was run on a test set of 7000 targets (see Materials and Methods for details). The first column represents the run performed with default settings, and in each subsequent column one variable is modified (see Table 3 for details).

**Table 2.**
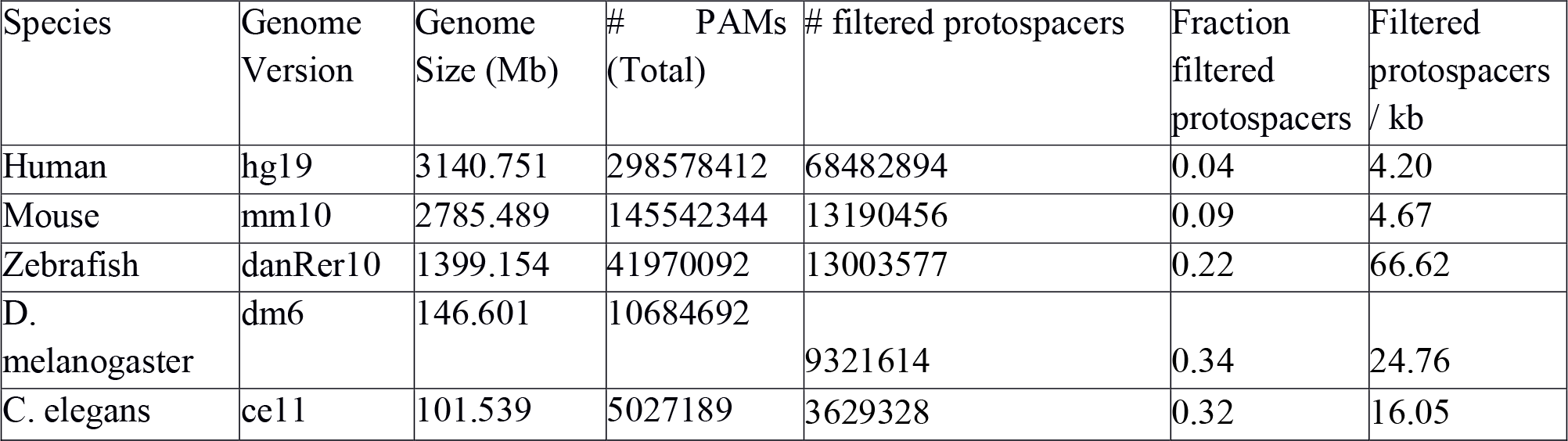
Species analysed by CRISPETa and for which off-target databases were compiled. Filtered protospacers are those passing default off-target and efficiency score cutoffs.

To estimate efficiency, candidate protospacers are scored using the logistic regression measure of Doench et al (5). This model was trained on experimental assays for 6085 and 1151 sgRNAs tiled across six mouse and three human genes, respectively. This score predicts sgRNA efficiency based on informative nucleotide preferences both within the core 20mer and in its immediate flanking nucleotides. These nucleotide preferences are summarised in Figure 1C. Protospacers passing defined off-target and on-target thresholds are retained-henceforth referred to as “filtered protospacers”. In contrast to off-target filtering, global efficiency scores are more consistent across genomes, removing from 60-70% of protospacers in the five genomes tested (Figure 1C and Table 2). Together, off-target and efficiency score filters eliminate 96% of candidate protospacers in human. There is markedly lower density of usable filtered protospacer sequences in vertebrates compared to invertebrates (Figure 2A).

In the final step, optimal sgRNA pairs are selected. First, all possible pairs of filtered protospacers are enumerated and ranked. Two ranking approaches are available: by combined efficiency score (default), or by length of deleted region. Ranking by score will tend to result in pairs that are more evenly distributed throughout the targeting region, but with a heterogeneous distribution of deletion sizes. Conversely, ranking by length favours shorter intervals within the constraints of the targeting design. Short segments may be more efficiently deleted (26), but will tend to be clustered into a smaller genomic region.

The top ranked pairs, up to a user defined maximum of n, are returned for each feature. In principle, a single high-scoring sgRNA may end up contributing to many or all of the highest-scoring pairs. To control this process, the “diversity” measure is used to control the maximum fraction of pairs containing a single sgRNA sequence (Table 1).

Finally, the user may specify constraints in sgRNA pair selection based on the plasmid construction method. Many plasmids employ the U6 promoter, which requires the sgRNA sequence to commence with a “G”. For instance, the DECKO plasmid expresses two sgRNAs in tandem from U6 and H1 promoters, thus requiring the 5’ sgRNA to commence with G (1). The “construction method” variable allows users to incorporate this constraint.

CRISPETa returns a ranked series of paired sgRNA constructs for each target. Sequences are output in FASTA format suitable for immediate ordering from commercial oligonucleotide synthesis services. Summary statistics and figures are produced for each design job.

### Performance of CRISPETa and effect of changes in parameters

We tested the standalone pipeline using a set of 7000 target genomic features compiled from a mixture of sources (see Materials and Methods). At default settings, CRISPETa returns successful, full depth (n=10) designs for 70% of features, with a further 17% of intermediate depth (0<n<10) and 13% failures (Figure 2C, Table 3). Performed on a workstation with CentOS6, 86.6 Gb of memory and 12 CPUs (Intel(R) Xeon(R) CPU E5649 @ 2.53GHz), the analysis took 44 minutes with a maximum RAM requirement of <100 MB.

**Table 3.** Summary of benchmarking results. Analyses were performed on a set of 7000 regions composed of different human target types (see Methods for details). % full depth refers to the percent of targets receiving 10 sgRNA pair designs. Designed targets refers to the total number of target features receiving full or partial depth designs.

| Name | Non-default parameter | Wallclock Time (s) | % full depth | % partial depth | Mean pairs per target | Mean protospacer score | Mean pair score | Mean pair distance | Total sgRNA pairs | # Input targets | # Designed targets |
| --- | --- | --- | --- | --- | --- | --- | --- | --- | --- | --- | --- |
| Default |  | 2639 | 70 | 17 | 8 | 0.52 | 1.05 | 1373 | 54441 | 7000 | 6058 |
| Diversity0.1 | v=0.1 | 4748 | 0 | 87 | 1 | 0.63 | 1.26 | 1372 | 6058 | 7000 | 6058 |
| Diversity 1 | v=1 | 4764 | 71 | 15 | 9 | 0.53 | 1.06 | 1373 | 55330 | 7000 | 6058 |
| off1 | t=1,0,1, x,x | 4869 | 84 | 8 | 9 | 0.58 | 1.16 | 1451 | 61378 | 7000 | 6444 |
| off2 | t=1,0,2, x,x | 4783 | 87 | 6 | 9 | 0.6 | 1.2 | 1449 | 63083 | 7000 | 6544 |
| off3 | t=1,0,3, x,x | 4802 | 89 | 5 | 9 | 0.61 | 1.22 | 1449 | 63832 | 7000 | 6584 |
| off4 | t=1,0,4, x,x | 4827 | 89 | 5 | 9 | 0.61 | 1.22 | 1448 | 64252 | 7000 | 6613 |
| off5_1 | t=1,1,5, x,x | 4883 | 91 | 4 | 9 | 0.62 | 1.24 | 1447 | 65173 | 7000 | 6670 |
| pScore0.6 | sp=0.6 | 4878 | 64 | 21 | 8 | 0.54 | 1.07 | 1367 | 52333 | 7000 | 6012 |
| pScore1 | sp=1 | 4691 | 31 | 40 | 5 | 0.6 | 1.21 | 1325 | 34196 | 7000 | 4978 |
| pScore1.2 | sp=1.2 | 5318 | 11 | 42 | 3 | 0.67 | 1.33 | 1243 | 18793 | 7000 | 3706 |
| Pscore1.6 | sp=1.6 | 4658 | 0 | 8 | 0 | 0.83 | 1.66 | 1109 | 1000 | 7000 | 569 |
| design_2000 | du=2000<br>dd=2000 | 9266 | 95 | 2 | 9 | 0.70 | 1.40 | 2872 | 67313 | 7000 | 6786 |

This benchmarking was repeated several times, in each case modifying a single parameter (Figure 2C and Table 3). As expected, strengthening the diversity requirement resulted in a drastic reduction of design success (“diversity0”), while a complete relaxation (“diversity1”) did not produce a substantial gain. Some improvement was observed when relaxing off-targeting, but this benefit is negligible after “off1” (allowing a single other match with two mismatches, (0:1, 1:0, 2:1, 3:x, 4:x). As expected, increasing the paired score threshold has a strong effect on design depth, particularly after 0.6 (“pScore0.6”) (default 0.4). The most dramatic improvement was observed when the length of the design region was increased to 2000 bp, boosting the fraction of successfully targeted regions from 70% to 95%. Thus, by adjusting these parameters, the depth of library designs can be optimised for each target set.

### Genome-scale deletion libraries for non-coding genomic elements in five species

We next used CRISPETa to design knockout libraries for a variety of genomic element classes that cannot be targeted by traditional RNAi’ either because their function is not thought to depend on RNA production (eg ultraconserved elements [UCEs])(21)’ or because their RNA product is too short (eg microRNAs) (see Table 4). We also created a collection of 3170 random intergenic target regions in human as a reference and for use as negative controls in screening projects. An example is shown in Figure 3A’ created using the standard output of CRISPETa’ where the IRX3 gene promoter and an upstream ultraconserved element (UCE) are targeted.

**Figure 3:**
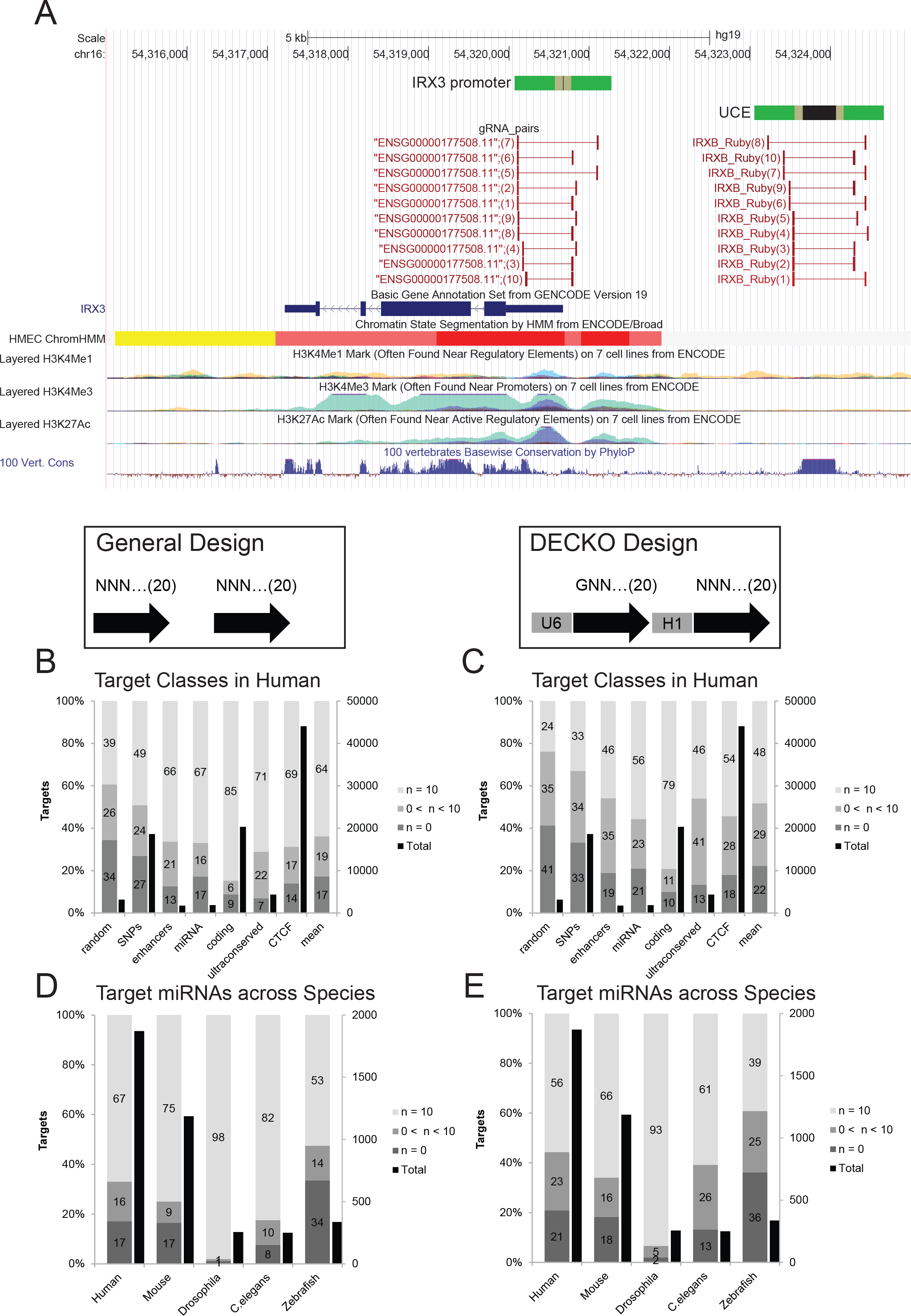
Genome-wide knockout libraries for classes of genomic elements. (A) An example of paired sgRNAs designed against the upstream ultraconserved element (UCE) and promoter of the human *IRX3* gene. *IRX3* lies on the antisense strand. The exact target regions are shown in black, flanked by the design regions in green. The ten sgRNA pairs for each are denoted by red bars. Integrated chromatin marks from the ENCODE project (18) are displayed below, in addition to PhyloP multispecies conservation scores (30). Note the region of elevated conservation corresponding to the UCE. (B-E) Summary of paired sgRNA designs targeting entire classes of genomic elements. In each figure, the left scale and grey bars represent the design performance, as in Figure 2. The right scale and black bars indicate the total number of elements in each class. B&C show a series of genomic element classes for human, while D&E show designs for the entire set of annotated microRNA genes in five species. B&D designs were created with default settings, while C&E employed the “DECKO” filter, requiring one protospacer to commence with a “G” nucleoside to be compatible with the U6 promoter.

**Table 4.**
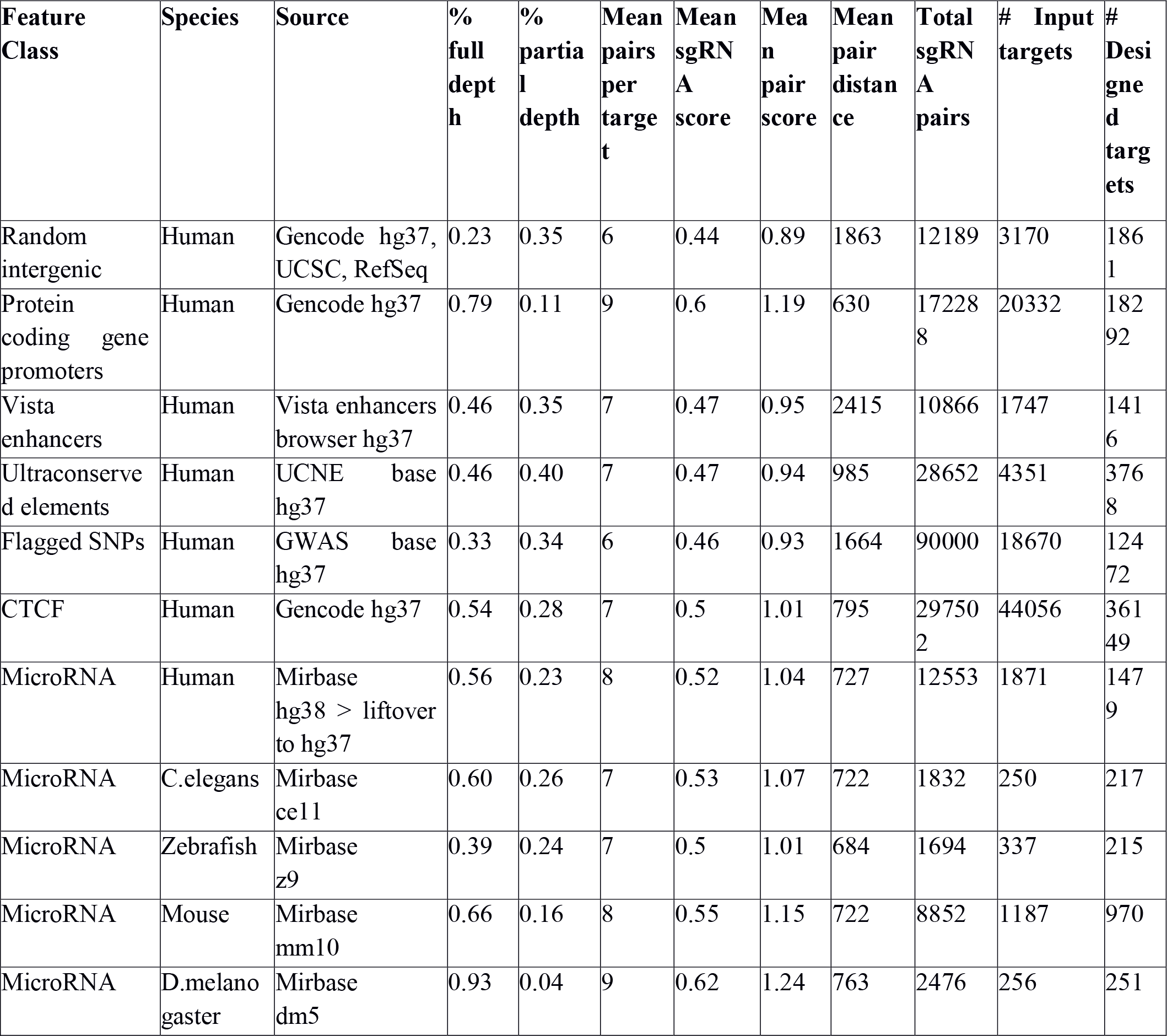
Pre-generated paired CRISPR libraries.

The characteristics of these libraries are shown in Figure 3B–E and Table 4. Overall’ 68% of features could be targeted at full depth’ with an additional 18% at incomplete depth. We observe considerable heterogeneity in the design success across classes’; with protein-coding gene promoters reaching a full depth for 85% of cases’; compared to 39% in random intergenic regions. We expect these differences arise from sequence uniqueness (affecting off-target frequency) and GC-richness (affecting PAM density and predicted efficiency). We created analogous targeting designs’; but imposing the requirement for DECKO-compatible sgRNA pairs’; and observed a considerable reduction in targeting depth (Figure 3B,C). This indicates that replacing the U6 promoter in the delivery vector could result in higher sgRNA coverage.

To compare performance across species’; we created designs targeting the entire annotated catalogue of microRNA genes in human’ mouse’ zebrafish’ D. *melanogaster* and C. *elegans* (Figure 3D,E). We observe considerably more efficient designs in non-mammalian species’; likely reflecting their more compact’ less repetitive nature. Nevertheless’; at default settings we managed to create full depth designs for 67% of human miRNA precursors’; and this could likely be improved by altering design parameters.

The entire sets of designs are available for download from the www.crispeta.crg.eu. Overall these results demonstrate the practicality of creating large-scale paired sgRNA knockout designs across diverse genomic element classes.

### Updated experimental methods for streamlined, economical and quantitative CRISPR seletion

To assess the efficiency of CRISPETa predictions’; we created two new experimental methods for CRISPR deletion. First’ we implemented a more streamlined and economical version of the recently-published DECKO method (1)’ called DECKO2 (Figure 4A–C). DECKO is a two-step cloning method for the creation of lentiviral vectors expressing a pair of sgRNAs. The backbone vector is compatible with lentivirus production and carries both puromycin resistance and fluorescent mCherry markers (Figure 4A). In the original protocol’ pDECKO is cloned from a single 165 bp starting oligonucleotide (“Insert-1”)’ which is compatible with both individual and complex library cloning.

**Figure 4:**
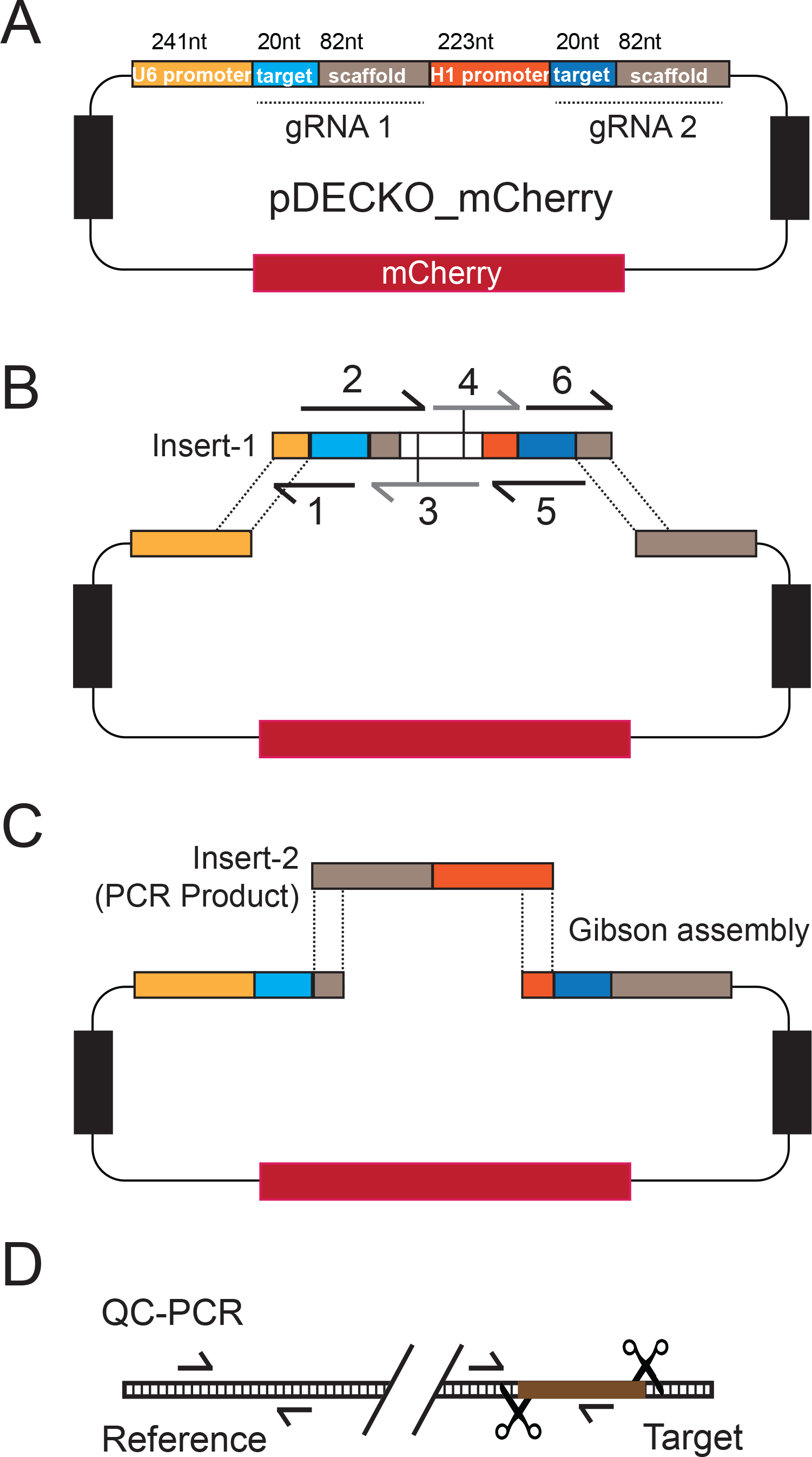
Updated experimental tools for genomic deletion and assaying its efficiency. (A-C) DECKO2, a streamlined and low-cost experimental protocol for generating paired sgRNA deletion lentiviral vectors. (A) The structure of the final pDECKO_mCherry vector (1). Blue indicates the variable regions of each sgRNA, corresponding to sgRNA recognition sites. Grey indicates constant scaffold regions. (B) Step 1 of the DECKO2 protocol: Insert-1 is simultaneously assembled from 6 overlapping oligonucleotides and inserted into the linearised backbone by Gibson assembly. Oligos 3 and 4 do not vary between targets. The result is the “intermediate” plasmid. Vertical bars in Insert-1 represent BsmbI restriction sites. (C) Step 2: The Insert-2 sequence, amplified by PCR, is inserted into linearised intermediate plasmid by Gibson assembly. (D) Outline of the QC-PCR method for assessing deletion efficiency. Concentration of unmutated’ wild-type target sites is normalised to the reference amplicon’ to control for template gDNA concentration.

DECKO2 is optimised for cloning individual CRISPR targeting constructs due to two novelties: first’ Insert-1 is assembled from a series of six shorter’ overlapping oligonucleotides (two of which are invariant for all targets)’ thereby reducing oligonucleotide synthesis costs by ~70% (Figure 4B and Supplementary File 5). Second’ conventional ligation is eliminated’ with both cloning steps relying on efficient and robust Gibson assembly (27). We did not observe any resulting decrease in cloning efficiency. To facilitate cell sorting’ we tested a number of combinations of Cas9 with fluorescent markers’; all of which displayed similar deletion efficiency (Supplementary File 6). We performed subsequent experiments in HEK293T stably expressing a simple Cas9-BFP fusion. This new protocol is described in the Materials and Methods and in more detail in Supplementary File 1.

In a second method, “quantitative CRISPR PCR” (QC-PCR), we evaluate CRISPR deletion efficiency in bulk (unsorted) cell samples (Figure 4D). The fraction of intact, wild-type target sites in a mixture of genomic DNA is quantified in real-time PCR, using primers amplifying the knockout region. Normalisation is performed using cells treated with non-targeting (EGFP) sgRNAs, and specificity is ensured using control primers amplifying a non-targeted region. QC-PCR can thus be used to quantify and compare the deletion efficiency of CRISPETa designs, by measuring the reduction of target sites in a mixed cell population.

### Quantitative experimental validation of CRISPETa

To test the effectiveness of sgRNA pairs designed by CRISPETa, we focussed on the human *MALAT1* locus. This lncRNA is a potent oncogene, which we previously silenced using DECKO (1,28). This time, we chose to delete two regions: a conserved upstream element with enhancer-like chromatin modifications (“enhancer”) and a region of conserved exonic sequence (“exon”) (Figure 5A). For each one, we used CRISPETa to design sgRNA pairs, and selected the three highest scoring pairs and one lower scoring pair (details can be found in Supplementary File 7). HEK293T Cas9-BFP cells were transfected with pDECKO, and selected by antibiotic resistance for 6 days, after which their genomic DNA (gDNA) was extracted (Figure 5B).

**Figure 5:**
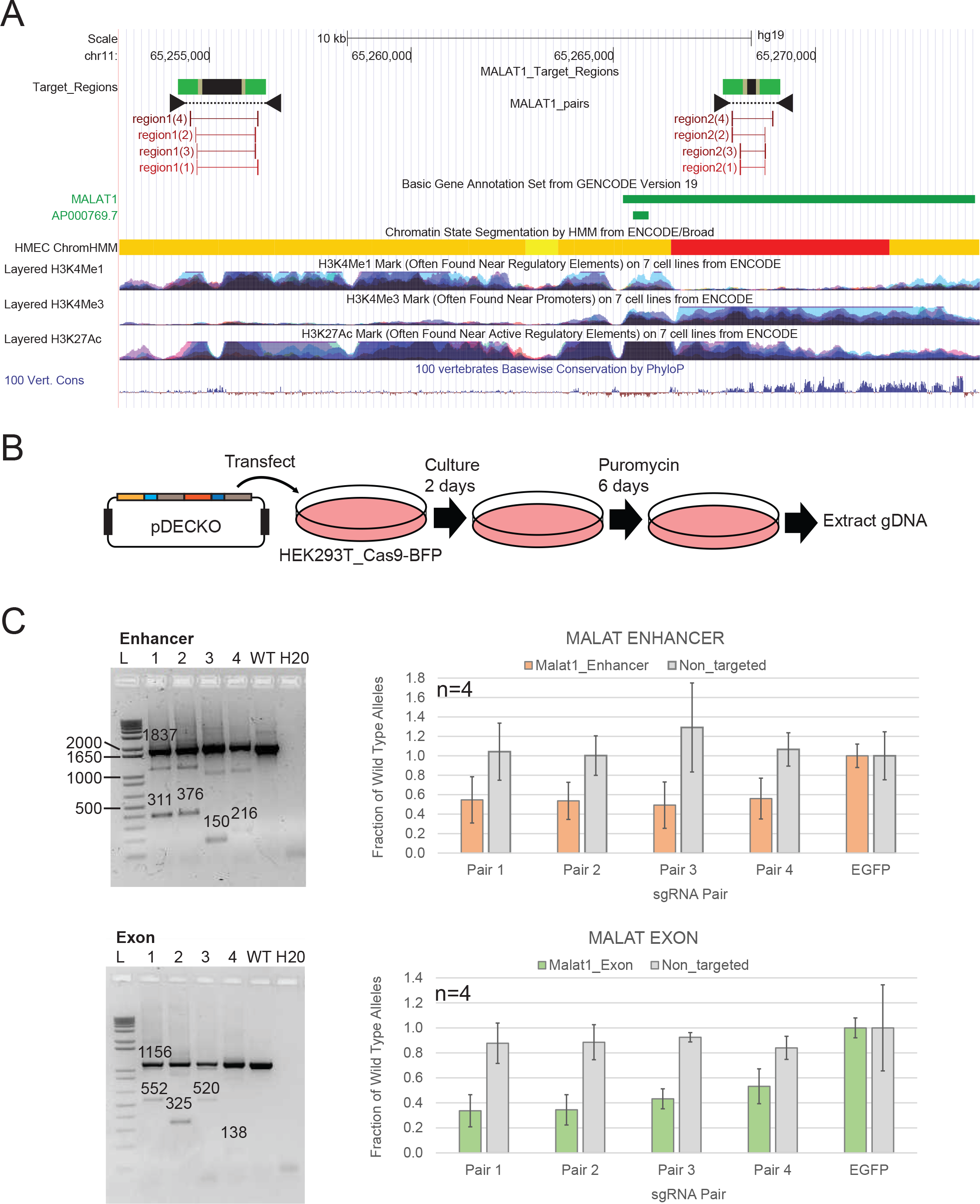
Experimental validation of paired sgRNA designs. (A) The human *MALAT1* locus. The *MALAT1* lncRNA gene’ shown in green’ lies on the positive strand. The two selected target regions are shown: the conserved upstream enhancer-like region (note the overlap with H3K4Me1 and H3K28Ac modifications)’ and the exonic region. As before’ target regions are shown in black’ and sgRNA design regions in green. sgRNA pairs used below are represented by red bars’; and genotyping primers as black arrowheads. (B) Overview of the experimental scheme. (C) Results for Enhancer region (upper panel) and Exon region (lower panel). Left: Agarose gel showing genotyping results from bulk (unsorted) cells’; with primers flanking the deleted region. 1-4 designate sgRNA pairs. “WT” indicates control cells transfected with pDECKO targeting the EGFP sequence. Numbers on the gel refer to the expected size for the PCR amplicons. Right: QC-PCR results from four independent biological replicates. Y-axis shows the normalised fraction of unmutated’ wild-type alleles’; using primers amplifying the targeted region (red/green)’ or a distal’ non-targeted region (grey). Data were normalised to control cells transfected with pDECKO_EGFP. Error bars denote the standard deviation of four independent biological replicates. The differences between treated cells and control cells were statistically significant for all four sgRNA pairs’; for both target regions (P>0.01’ paired *t* test).

Conventional PCR genotyping of transfected cells’; DNA showed amplification products consistent with target site deletion for all pDECKO constructs, but not for control cells (Figure 5C, left panels). QC-PCR analysis of independent biological replicates showed loss of ~40% of enhancer target sites for each of the four sgRNA pair designs (Figure 5C, right panels). A non-targeted genomic region was not affected (“Non-targeted”). Higher efficiencies were observed for the exon-targeting constructs, yielding >60% efficiency for the top two sgRNA pairs. We did not observe a strong difference in the deletion efficiency between the four sgRNA pairs targeting the enhancer, although for the exon region, the lower-scoring two constructs displayed reduced efficiency. This underlines the value of using predicted efficiency scores in sgRNA selection, and supports the effectiveness of CRISPETa-predicted sgRNA pairs.

### Mutant RNA arising from genomic mutation

We next sought to verify that the engineered deletions in the *MALAT1* exon result in the expected changes to transcribed RNA. cDNA was generated from bulk cells treated with pDECKO vectors targeting MALAT1 exon. Given that cells were not selected, this sample should contain a mixture of RNA from both wild-type and mutated alleles. RT-PCR using primers flanking the targeted region amplified two distinct products, of sizes expected for wild-type and deleted sequence (Figure 6). The specificity of these PCR products was further verified by Sanger sequencing. Therefore, targeted deletions by CRISPETa are reproduced in the transcriptome, and may be used in future dissect RNA functional elements.

**Figure 6:**
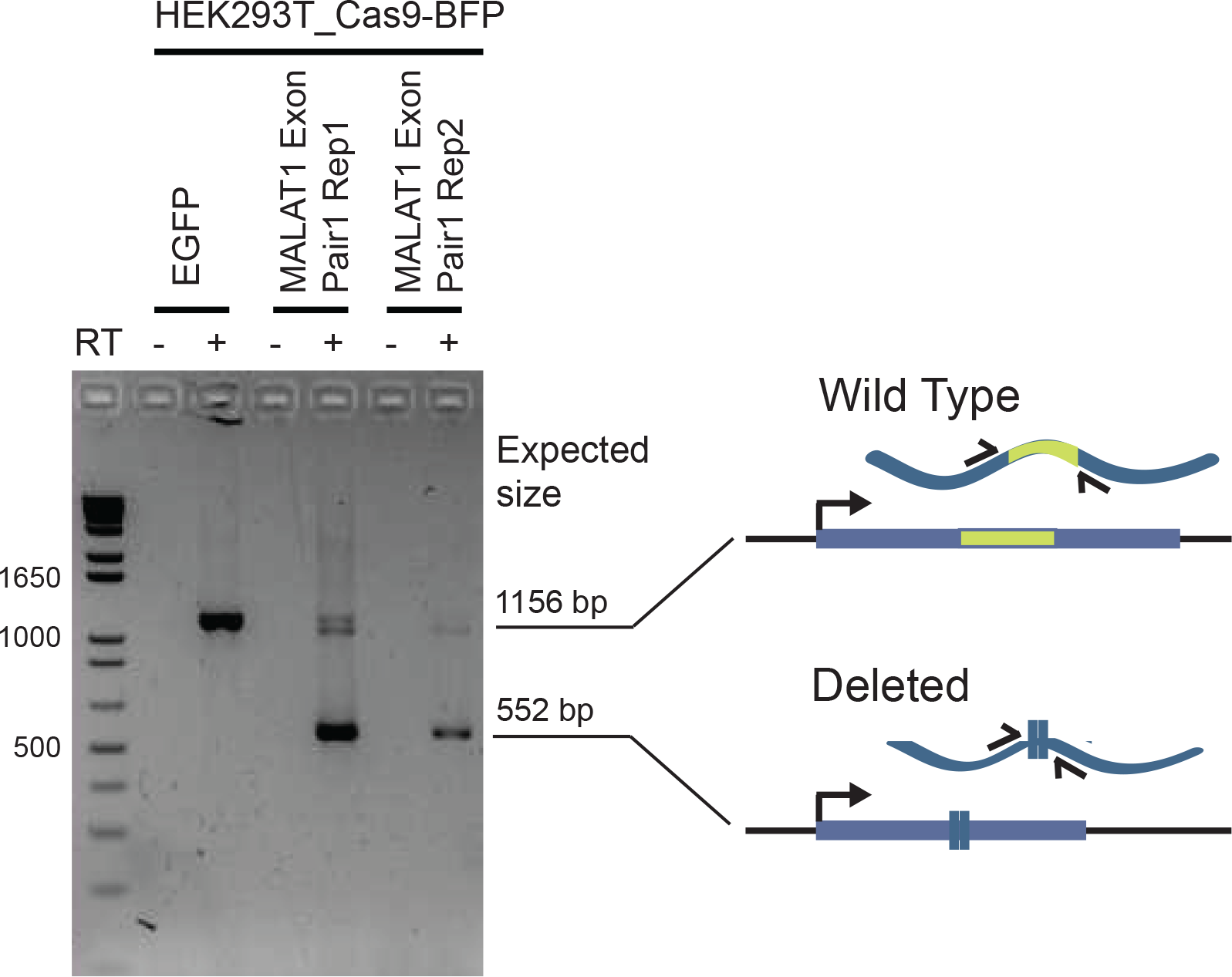
Observation of mutated MALAT1 RNA. RT-PCR was performed on RNA from bulk cells where MALAT1 exon region was deleted (sgRNA Pair 1’ in two biological replicates)’ or control cells transfected with pDECKO_EGFP. Primers flanking the deleted region were used’ and are expected to amplify fragments of the indicated sizes’; depending on whether the RNA arises from a wild type or a deleted allele. Specificity was ensured by the exclusion of the reverse transcriptase enzyme in control reactions (“RT-”).

### Genomic deletion under screening conditions at low multiplicity of infection

In future, the DECKO_mCherry and CRISPETa tools may be used in library screening-type experiments (4). The above experiments were carried out by transfection, meaning that each cell receives multiple copies of pDECKO plasmid. In contrast, library screening requires the integration of a single targeting sequence in each cell. This requirement is met at low multiplicity of infection (MOI) when ≤20% of cells are infected, a condition that can be conveniently monitored by mCherry fluorescence (29). Thus, in a final experiment we sought to test whether, under such conditions of single-copy integration, pDECKO_mCherry remains effective in genomic deletion.

For these experiments, we selected IMR90 fibroblasts since they are not transformed and hence more suitable model for phenotypic screening. Using a previously-validated pDECKO_mCherry lentivirus targeting the promoter of the *TFRC* gene (1), we infected cells at decreasing titres. By means of flow cytometry gated on mCherry fluorescence, we could monitor infection rate. Infected cells were cultured under antibiotic selection to create a pure population, then genotyped by PCR (Figure 7). We observed the presence of correctly mutated alleles in cells carrying single-copy pDECKO insertions. Thus, CRISPETa-designed libraries are likely to be suitable for pooled screens.

**Figure 7:**
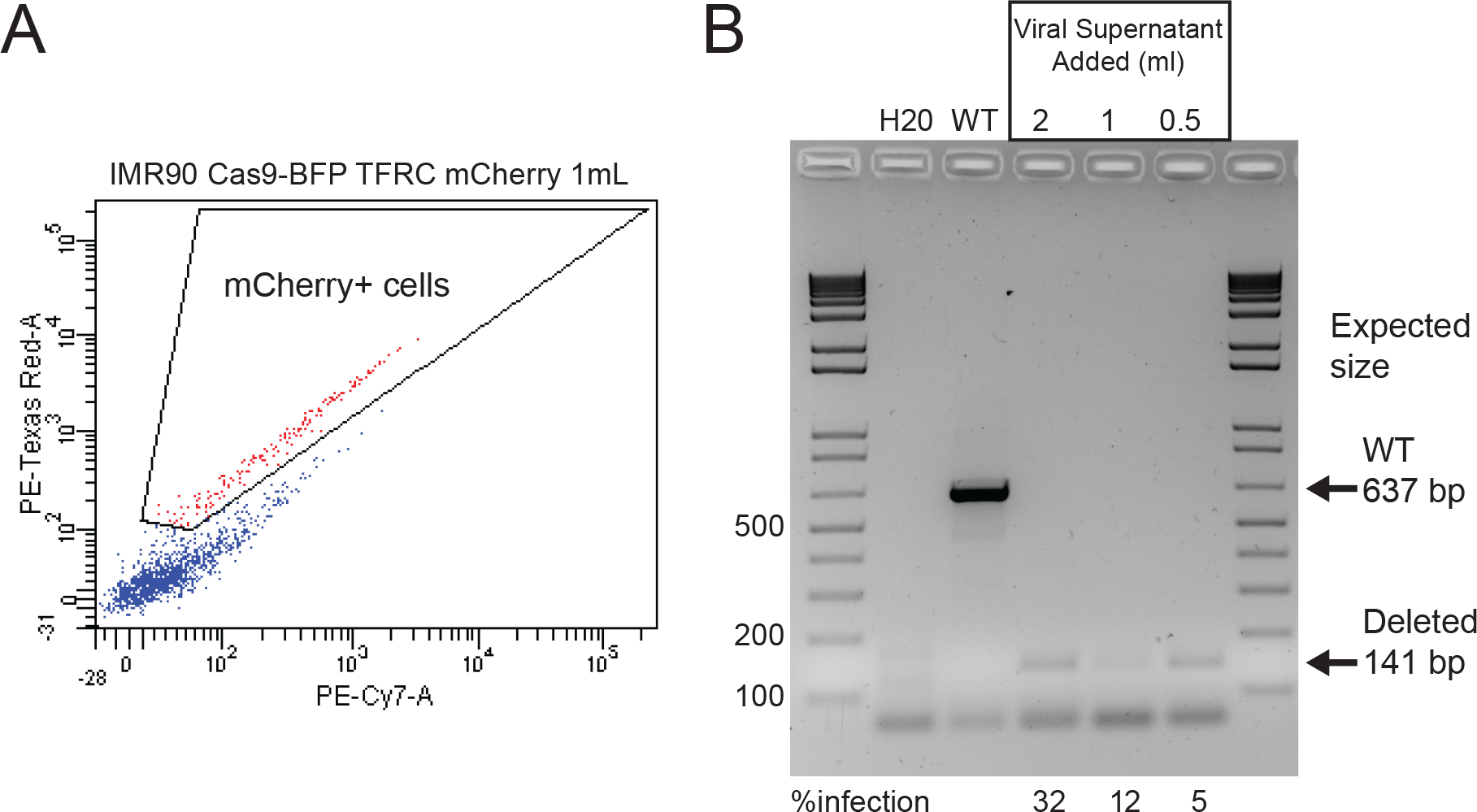
Efficacy of DECKO in screening conditions at low multiplicity of infection. (A) IMR90 cells stably expressing Cas9-BFP were infected with varying volumes of lentiviral supernatant carrying a pDECKO construct targeting the *TFRC* gene (TFRC_B construct’ see (1)). Infection rates were estimated by flow cytometry monitoring fluorescence from the mCherry gene carried by pDECKO. In this example’ infection rate is ~12% (see next panel). (B) Genotyping PCR carried out on genomic DNA from lentivirally-infected cells. The infection rate in the original sample is indicated below. Note that antibiotic selection was subsequently used to remove non-infected cells. Primers flank the deletion site’ and expected amplicon sizes for wild type (unmutated’ “WT”) and deleted alleles are indicated.

## Conclusions

We have here presented a versatile and scalable design solution for CRISPR deletion projects. To our knowledge’ CRISPETa is the first tool for selection of optimal sgRNA pairs. A key feature is its scalability’ making it equally suitable for focussed projects involving single target regions’; and screening projects involving thousands of targets. The user has a large degree of control over the design process. On-target efficiency is predicted using the latest’ experimentally-informed design algorithm’ while running speed is boosted by an efficient off-target calculation.

In the course of this work we developed an updated lentiviral CRISPR deletion tool. Compared to the original version’ DECKO2 represents a more cost-effective method for individual target knockout. The series of short oligonucleotides required cost less to synthesise from commercial vendors compared to a single 165 nt sequence employed previously (1). DECKO2 has also replaced the second ligation step of the original DECKO by Gibson assembly’ further simplifying the protocol (27).

The QC-PCR technique presented here now allows one to quantify and compare the efficiency of CRISPETa designs. For the 8 sgRNA pairs in two regions that we tested’ deletion efficiencies of ~40-60% were consistently observed. The induced deletions’; when occurring within a transcribed region’ are also observed in expressed RNA molecules. The suitability of this approach for screening is demonstrated by the fact that single-copy genomic insertions give rise to deletion of target regions.

CRISPR enables us to study the function of non-coding genomic elements in their endogenous cellular context for the first time. The power of CRISPR lies both in its versatility’ but also in its ready adaptation to large-scale screening approaches. The CRISPETa pipeline and experimental methods described here will’ we hope’ be useful for such studies.

## Acknowledgements

We thank members of the Guigo lab and the CRG Bioinformatics and Genomics Programme for many ideas and discussions, in addition to Carlo Carolis of the CRG Biomolecular Screening & Protein Technologies Unit. We thanks John G. Doench and David E. Root (Broad Institute of MIT) for generous advice regarding the implementation of sgRNA efficiency predictions. We acknowledge the administrative support of Romina Garrido. We also acknowledge support of the Spanish Ministry of Economy and Competitiveness, ‘Centro de Excelencia Severo Ochoa 2013-2017’, SEV-2012-0208. This work was financially supported by the following grants: CSD2007-00050 from the Spanish Ministry of Science, grant SGR-1430 from the Catalan Government, grant ERC-2011-AdG-294653-RNA-MAPS from the European Community financial support under the FP7 and grant R01MH101814 by the National Human Genome Research Institute of the National Institutes of Health, to RG. Ramon y Cajal RYC-2011-08851 and Plan Nacional BIO2011-27220 to RJ.

---

Supplementary File 1: Extended DECKO2 cloning protocol.

Supplementary File 2: Design spreadsheet for creating DECKO2 oligonucleotides.

Supplementary File 3: Estimation of QC-PCR primer efficiencies.

Supplementary File 4: Oligonucleotide sequences.

Supplementary File 5: Detailed figure of 6-oligo Insert-1 cloning.

Supplementary File 6: Comparing efficiency of fluorescent Cas9 variants.

Supplementary File 7: Details of *MALAT1* sgRNA pairs.

